# Natural speech re-synthesis from direct cortical recordings using a pre-trained encoder-decoder framework

**DOI:** 10.1101/2024.12.16.628596

**Authors:** Jiawei Li, Chunxu Guo, Edward F. Chang, Yuanning Li

## Abstract

Reconstructing perceived speech stimuli from neural recordings is not only advancing the understanding of the neural coding underlying speech processing but also an important building block for brain-computer interfaces and neuroprosthetics. However, previous attempts to directly re-synthesize speech from neural decoding suffer from low re-synthesis quality. With the limited neural data and complex speech representation space, it is hard to build decoding model that directly map neural signal into high-fidelity speech. In this work, we proposed a pre-trained encoder-decoder framework to address these problems. We recorded high-density electrocorticography (ECoG) signals when participants listening to natural speech. We built a pre-trained speech re-synthesizing network that consists of a context-dependent speech encoding network and a generative adversarial network (GAN) for high-fidelity speech synthesis. This model was pre-trained on a large naturalistic speech corpus and can extract critical features for speech re-synthesize. We then built a light-weight neural decoding network that mapped the ECoG signal into the latent space of the pre-trained network, and used the GAN decoder to synthesize natural speech. Using only 20 minutes of intracranial neural data, our neural-driven speech re-synthesis model demonstrated promising performance, with phoneme error rate (PER) at 28.6%, and human listeners were able to recognize 71.6% of the words in the re-synthesized speech. This work demonstrates the feasibility of using pre-trained self-supervised model and feature alignment to build efficient neural-to-speech decoding model.

## Introduction

Speech language perception is a process in human auditory cortex that transforms continuous auditory stimuli of speech language into the perception of different levels of linguistic units such as phonemes, syllables, words, etc.[1] There are two main perspectives to study how these different levels of information processing and representation happen in the brain. One is neural encoding, where different feature representations of the stimuli are used to predict the neural responses during speech processing [2-6]; the other is neural decoding, where neural signals are used to decode and reconstruct features from the stimuli. Speech re-synthesis from cortical response is a method to study speech language perception by decoding the auditory stimuli from recorded neural activities. By evaluating the amount of information retrieved from the neural activity, we learn the content of the neural coding in certain cortical network during the process. This can be seen as a data-driven way to learn about neural coding. Furthermore, re-synthesizing or decoding continuous speech is also a critical algorithmic component in brain-computer interfaces, which directly transfer neural signal into speech and language [7-11]. However, brain-to-speech synthesizing is a more challenging task, compared to brain-to-text decoding. There is richer acoustic-phonetic and intonational information represented in the continuous speech space, compared to the discrete space of linguistic units like phonemes and words.

The first challenge of re-synthesizing continuous natural speech is experimentally accessing the corresponding neural activity, accounting for the highly dynamic nature of natural speech. Synthesizing natural speech requires not only veridical re-synthesis of lower-level acoustic features or discrete phonetic/linguistic labels, but also recovering the continuous sentential-level suprasegmental features and dynamics. Previous studies have approached brain-to-speech decoding and synthesizing, demonstrating the feasibility of decoding different levels of linguistic information from neural responses [12-14]. Given the limitations in spatiotemporal resolution and signal-to-noise ratio in non-invasive methods, it is more appealing to seek for more invasive methods for more robust and precise brain-to-speech synthesizing. Recent invasive electrophysiological methods have offered a deeper insight into the fine grain coding of different levels of information in the speech auditory cortex. Studies using invasive methods have shown that different spectral, temporal, acoustic, phonetic and formant [15], spectrogram [8, 16], phoneme [16], word [17, 18], word identity [19], speaker identity [7] can be decoded from brain recordings.

The second major challenge is the computational difficulty in mapping brain activity into continuous speech. Despite the high spatiotemporal resolution and the signal-to-noise ratio offered by invasive neural recordings, to overcome the highly nonlinear mapping between brain activity and speech language and to achieve high-performance in brain-to-speech/text decoding would still require intensive amount of training data. For example, it takes 17 days of chronic intracranial recording data to train recurrent neural networks that decode brain activity into naturalistic speech at a phoneme error rate of 25% [9]. In most clinical settings, such as during awake neurosurgeries or epileptic monitoring, it is expensive or infeasible to chronically record large amount of intracranial for speech tasks in specific. Compared to the state-of-the-art deep end-to-end speech neural networks [20, 21], the amount of data cumulated in a single neural speech dataset is orders of magnitude smaller than the typical size of speech dataset required to train complicated deep neural networks. Therefore, decoding natural speech from intracranial recordings poses unique challenges from both data and algorithmic perspectives. How to build efficient and high-performance decoding algorithm that operates on a limited amount of training data is critical **for the development of brain-to-speech decoding techniques**.

Recent studies on pre-trained self-supervised models have offered critical insight into these questions. There is a growing body of literature identifying representational similarity between pre-trained deep speech and language models and the human brain activities [22-24]. These findings suggest that the feature embeddings extracted from deep neural network models trained on large corpus of speech and text can be aligned to the corresponding neural responses, and there exist simple near-linear mappings between the representation spaces of artificial intelligence (AI) models and the brain. Furthermore, recent breakthroughs in deep generative models of speech and language have also made it possible that high-quality speech and text can be synthesized and generated from an embedding space trained from large corpus of unlabeled speech and text [25-27]. Putting these two perspectives together, it suggests a possible indirect brain-to-speech transformation where the brain activity is first aligned and mapped to the learned embedding space of a deep speech encoding model and then another deep generative model is used to generate naturalistic speech from the embedding space. The first part relies on the relatively simple mapping between brain and deep speech models, and can be trained with relatively small set of neural data; the second part is more data-hunger but can be trained purely on unlabeled speech data, which is easily accessible with low cost.

In this study, we designed a brain-to-speech decoding model following these strategies. We used high-density electrocorticography recordings to get high spatiotemporal resolution coverage of the cortical areas responsible for speech perception. Unlike previous brain-to-speech decoding methods that directly train end-to-end models from scratch, we trained a two-phase model: a light-weighted neural feature adaptor that mapped neural signal onto the latent space of deep speech encoder, and a pre-trained speech generator that generates high-fidelity natural speech sound from the speech encoder. We showed that our methods reliably reconstruct perceived natural speech, including phonemes, words, and temporal landmarks, using only 20 minutes of electrocorticography (ECoG) recordings as neural training data. This underscores the importance of incorporating pre-trained deep speech model in neural decoding applications.

## Results

### Overview of the proposed speech reconstruction method

Our proposed decoding model consists of two training stages: 1) self-supervised deep speech generator pre-training stage; 2) supervised neural feature adaptor training stage (**Fig. 1**).

**Fig 1.**
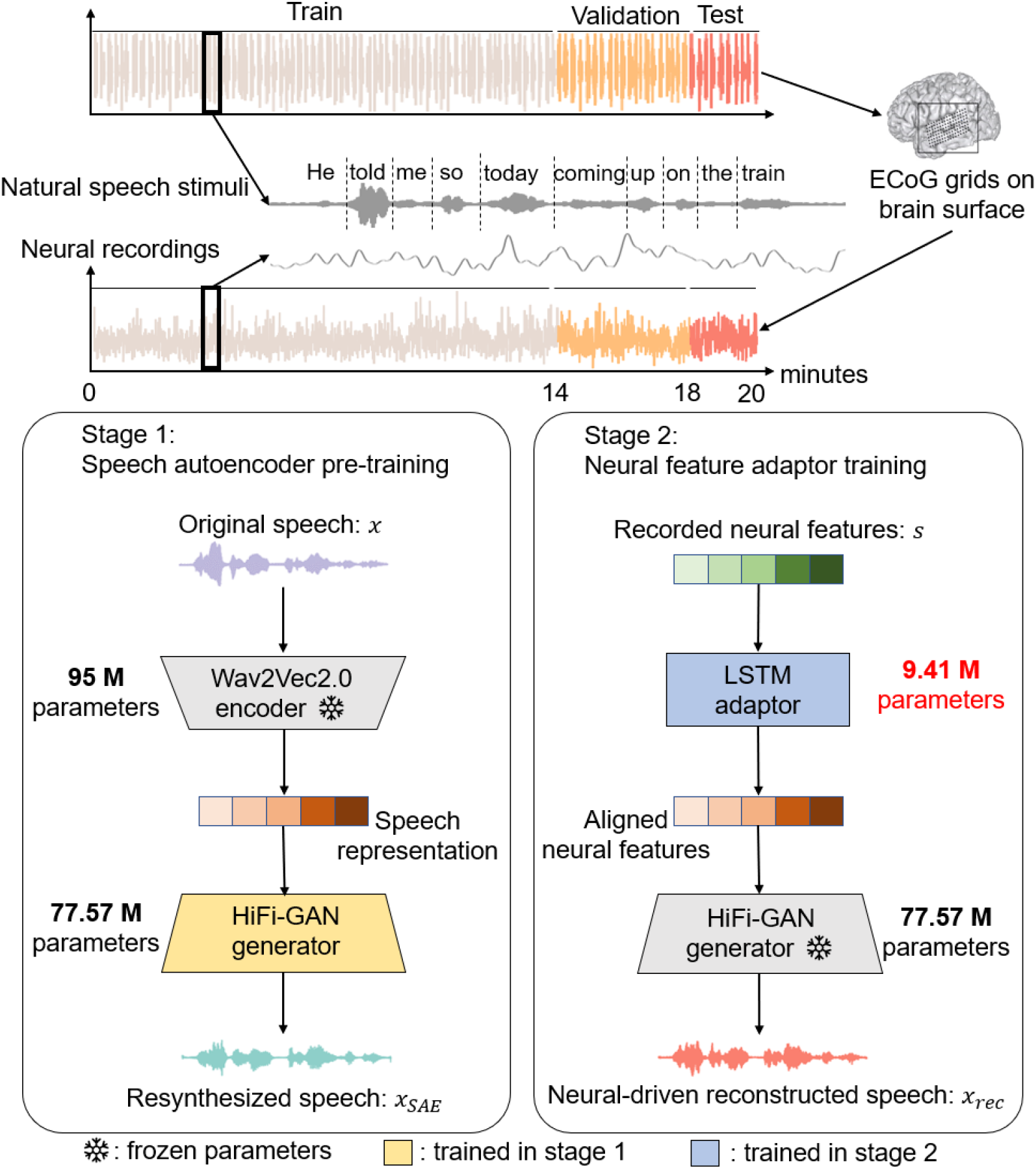
The proposed decoding model involves two training stages for high-fidelity speech synthesis. In Stage 1, a HiFi-GAN generator is pre-trained to synthesize natural speech from features extracted by the pre-trained Wav2Vec2.0 encoder, using a multi-receptive field fusion (MRF) module and adversarial training with discriminators to capture various speech patterns. This stage uses a large corpus (LibriSpeech) to enhance speech representation learning. In Stage 2, a lightweight neural feature adaptor, consisting of a three-layer bi-directional LSTM with 9.41M parameters, maps neural activity to speech representations. This adaptor aligns neural features (recorded ECoG signals) with speech features, allowing the HiFi-GAN generator to re-synthesize high-quality speech from neural data. Notably, we collected ECoG data from 9 monolingual native English participants as they each listened to English sentences from the TIMIT corpus, resulting in 20 minutes of neural recording data per participant. Furthermore, for each participant, 70%, 20%, and 10% of the recorded neural activities are respectively allocated to the training, validation, and test sets. The number of parameters in the LSTM-adaptor is much smaller (9.41M) compared to Wav2Vec2.0 (95M) and HiFi-GAN (77.57M). Additionally, a sentence excerpt has been extracted from the data for use as an example of a natural language sentence within the figure.

In the first stage, our goal is to train a deep speech generator to synthesize natural speech waveforms from latent features. To do so, we employed an encoder-decoder architecture speech Autoencoder (SAE), where the speech encoder transformed speech waveform into latent feature representations while the speech decoder (generator) resynthesized speech waveform from the latent representations (**Fig. 1**). As we argued in the previous section, a key design consideration was to use latent representations that were closely related to the human brain responses. Therefore, we took a pre-trained Wav2Vec2.0 [20] as the speech encoder, which is a Transformer-based unsupervised model that encodes speech waveform into a feature space correlated to human brain [23, 24]. For the generator part, we trained a HiFi-GAN generator[25]. HiFi-GAN is a generative adversarial network for high-fidelity speech synthesis. The HiFi-GAN generator and discriminators are adversarially trained to synthesize high-quality natural speech using speech features extracted by the pre-trained Wav2Vec 2.0 speech encoder. In this stage, the parameters of Wav2Vec2.0 speech encoder were frozen while the parameters of HiFi-GAN were tunable. This strategy enables us to overcome the limitations of neural recording data and implement a relatively large-scale deep speech generator (∼100M parameters) using pure unlabeled speech corpus.

In the second stage, our goal is to train a lightweight feature alignment model that transforms the neural activity to the latent speech representations. This facilitates the ultimate neural-to-speech synthesis of high-fidelity natural speech sentences through the deep speech generator pre-trained in the first stage. The linear correlations between the neural activity and the latent speech representations allow for the mapping of recorded neural signals onto latent speech features through a lightweight model. The adaptor comprises a three-layer bi-directional LSTM (Long-Short Term Memory) and notably contains a significantly smaller number of parameters (9.41M) in comparison to the Wav2Vec2.0 speech encoder (95M parameters) and HiFi-GAN decoder (77.57M parameters). In this stage, the parameters of HiFi-GAN generator were frozen while the parameters of the light-weight adaptor were tunable. As a result, adaptor model can be easily trained using a limited set of neural data. For each participant, 20 minutes of neural data were collected when they listened to natural speech. These data were split to train, validate and test the model.

### The pre-trained deep speech generator can reconstruct high-fidelity continuous natural speech

First, we evaluated the performance of the self-supervised deep speech Autoencoder built in stage 1. We trained the model on LibriSpeech, a 960-hour unlabeled speech corpus [28]. We then evaluated the performance on an independent test set of TIMIT [29], a labelled speech dataset annotated at the phoneme level. We evaluated the quality of the reconstructed speech using both subjective and objective metrics. To better benchmark the performance, we also generated surrogate speech from the original raw waveforms with additive noise at different signal-to-noise ratios, and evaluated the quality of these surrogate speech using the same metrics.

Objective assessment was conducted using extended short-time objective intelligibility (ESTOI[30]) and mel-spectrogram mean-square error (mel-spectrogram MSE). ESTOI, ranging from 0 (poor) to 1 (excellent), quantifies the intelligibility of speech synthesis by measuring distortion in spectrotemporal modulation patterns, sensitive to spectral and temporal inconsistencies [30]. The ESTOI score for the synthesized speech from our model is 0.825 ± 0.045 (**Fig. 2b**). For comparison, we also computed these metrics on surrogate speech with additive noise [31] (measured in dB). The quality of the reconstructed speech was better than adding -20dB babble noise to the raw original speech (ESTOI = 0.751 ± 0.038, t(49) =7.272, p=2.517×10^−9^, two-sided, **Fig. 2b**). The mel-spectrogram MSE measures the mel-spectrogram difference between the reconstruct speech and the ground truth. The speech sentences processed by our proposed model had mel-spectrogram MSE of 0.148 ± 0.017, which was comparable to the addition of -30dB additive noise to the raw speech (0.144 ± 0.014, t(49) = 0.212, p = 0.416, two-sided, **Fig. 2c**).

**Fig 2.**
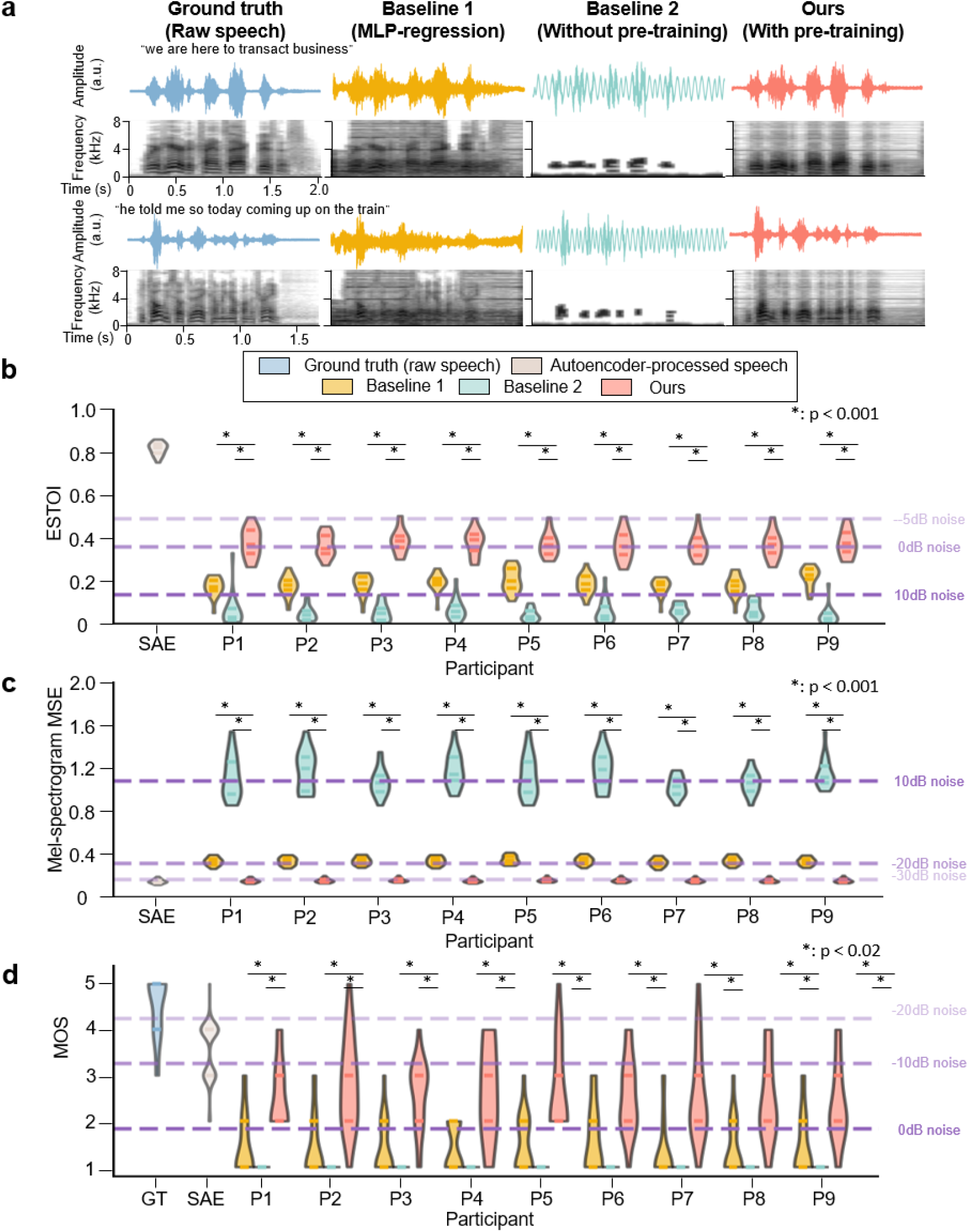
Comparative assessment of speech enhancement methodologies through subjective and objective measures of quality and intelligibility. a.Speech waveform and mel-spectrogram comparisons. Depicted are waveforms (time vs. amplitude) and mel-spectrograms (time vs. frequency ranging 0-8 kHz) for illustrative speech samples (Ground truth, MLP-reconstructed speech, scratch-trained speech and neural-adapted speech). b.Objective intelligibility assessment via ESTOI. The violin plots represent the distribution of ESTOI scores, an objective metric ranging from 0 (poor) to 1 (excellent), evaluating the preservation of spectrotemporal modulations in re-synthesized speech against the Ground Truth. Higher scores for our With Pre-training method, marked by asterisks, highlight its effectiveness in minimizing spectral and temporal distortions. * denotes that the speech re-synthesis with pre-training statistically significantly outperforms baseline methods in terms of voice quality. The three shades of purple dashed lines from light to dark represent the average ESTOI scores after adding babble noise at -5 dB, 0 dB, and 10 dB to the original speech waveform. SAE: Autoencoder-processed speech. P1-P9: participant 1 – participant 9. c.Objective evaluation using mel-spectrogram mean square error (MSE). The violin plot depicts the distribution of mel-spectrogram MSE scores, evaluating the fidelity of spectral representation in re-synthesized speech against the Ground Truth. Lower MSE values, particularly highlighted with asterisks for our With Pre-training method, indicate superior accuracy in capturing mel-spectrogram features compared to baseline methods. * denotes statistically significant improvements in spectral fidelity and robustness attributed to our approach. The three shades of purple dashed lines represent the average MSE scores after introducing babble noise at 10 dB, -20 dB, and -30 dB to the original speech waveform. SAE: Autoencoder-processed speech. P1-P9: participant 1 – participant 9. d.Subjective quality evaluation through mean opinion scores (MOS). Human evaluators rated both raw and re-synthesized speech samples on a scale from 1 (poor) to 5 (excellent), with the violin plot summarizing these MOS ratings across Ground truth, MLP-reconstructed speech, scratch-trained speech and neural-adapted speech. Asterisks indicate statistically significant enhancements in perceived intelligibility and quality attributed to our proposed method. * denotes that the speech re-synthesis with pre-training statistically significantly outperforms baseline methods in terms of voice quality. The three shades of purple dashed lines from light to dark represent the average MOS scores after adding babble noise at -20 dB, -10 dB, and 0 dB to the original speech waveform. GT: ground truth (raw speech). SAE: Autoencoder-processed speech. P1-P9: participant 1 – participant 9.

In addition to the objective metrics, subjective evaluation was performed using mean opinion score (MOS[32]). 20 human evaluators rated the speech intelligibility on a scale from 1 (poor) to 5 (excellent). The MOS for the re-synthesized speech reached 3.543±0.409, which was on par with the MOS of -10dB additive noise (MOS = 3.254 ± 0.741, t(49) = 1.441, p=0.078, two-sided t-test) on natural speech. For the original speech, the MOS was at 4.287 ± 0.321, which was comparable to -20dB additive noise (MOS = 4.234 ± 0.463, t(49) = 1.540, p = 0.065) (**Fig. 2c**, further details provided in the Methods section).

Our evaluations demonstrate that the self-supervised deep speech generator performs robustly, maintaining high intelligibility and quality. Both objective measures (ESTOI) and subjective assessments (MOS) indicate that the reconstructed speech is comparable to speech with moderate levels of noise, highlighting the effectiveness of our model in generating clear and intelligible speech from unlabeled data. This pre-trained model laid the foundation for our next steps of building neural-to-speech decoder.

### Evaluations of the acoustic quality of the neural-driven re-synthesized speech

Next, we built the neural-to-speech feature-adaptor and evaluated the performance of the neural-driven reconstructed speech. We collected ECoG data from 9 monolingual native English participants when they each listened to English sentences from TIMIT corpus. This yielded 20 minutes of neural recording data from each participant. This data was used for supervised training of the neural feature adaptor model in stage 2. We also used the same data to train two baseline models for comparison (**Table 1**). The first baseline model employed a three-layer multi-layer perceptron (MLP) as an end-to-end model that directly mapped neural recordings onto the mel-spectrogram without any pre-trained decoder. The second baseline model shared the same Hifi-GAN decoder and LSTM feature adaptor architecture with our model, but was trained merely on neural data from scratch, without any self-supervised pre-training.

**Table 1.** Summary of network architectures and training methods. The re-synthesized speech sentences from neural activity were compared by two baseline methods to demonstrate the necessity of encoder-decoder architecture and GAN decoder pre-training. The first baseline used a three-layer MLP to map neural recordings directly onto mel-spectrograms, demonstrating the limitations of direct synthesis without pre-trained decoders. The second baseline involved training a speech re-synthesis model from scratch without Librespeech pre-training, highlighting the significance of pre-training by comparing its performance to models incorporating this phase. The network architectures and training methods in the two baselines and our method are marked with a ‘+’ (used) or ‘ −’ (not used).

| Training methods | encoder-decoder architecture | GAN decoder pre-training | neural-driven supervision |
| --- | --- | --- | --- |
| Baseline 1 (End-to-end MLP-regression) | - | - | + |
| Baseline 2 (without pre-training) | + | - | + |
| Ours | + | + | + |

We evaluated the re-synthesized sentences for each participant in the test set. Similar to evaluate the performance of the deep speech generator in stage 1 (**Fig. 1**), the re-synthesized speech was assessed using both subjective and objective measures, including ESTOI score, mel-spectrogram MSE and MOS. The averaged ESTOI score **(Fig. 2b**) for the re-synthesized speech across participants was 0.377±0.013, with the best performance across participants achieving 0.384 ± 0.011. The ESTOI score of re-synthesized speech is comparable to natural speech with 0dB natural speech (ESTOI = 0.356 ± 0.012, t(179) = 1.297, p = 0.210, two-sided t-test). These results significantly outperformed the two baseline methods, with the average ESTOI scores for the MLP-reconstruction and scratch-training being 0.186±0.010 (t(179) = 9.150, p = 8.008×10^−11^, two-sided t-test) and 0.052±0.009 (t(179) = 14.546, p = 4.419 × 10^−17^, two-sided t-test), respectively. The best ESTOI performances across participants for these baseline methods were 0.219 ± 0.010 and 0.065 ± 0.010, respectively.

The average mel-spectrogram MSE (**Fig. 2c**) for the neural-driven re-synthesized speech across participants was 0.151±0.018, with the best performance across participants achieving 0.148 ± 0.017. The mel-spectrogram MSE of re-synthesized speech is comparable to natural speech with -30dB noise (mel-spectrogram MSE = 0.159 ± 0.028, t(198) = -1.353, p = 0.192, two-sided t-test). These results outperform the two baseline methods as well (MLP-reconstruction: MSE = 0.333 ± 0.037, t(179) = 80.946, p = 3.132 ×10^−143^, two-sided t-test; scratch-training: MSE = 1.120 ± 0.159, t(179) = 71.174, p = 1.573 ×10^−133^, two-sided t-test).

Furthermore, the average MOS (**Fig. 2d**) for the neural-driven re-synthesized speech across participants was 2.617 ± 0.196, with the best performance across participants reaching 2.800 ± 0.219. The MOS score of re-synthesized speech perform between natural speech with -10dB and 0dB noise (-10dB: MOS = 3.254 ± 0.741, t(79) = 8.876, p = 8.482×10^−14^, two-sided t-test; 0dB: MOS = 1.835 ± 0.595, t(79) = -6.081, p = 2.006 ×10^−8^, two-sided t-test).These results also outperformed the baseline methods, where the average MOS for the MLP-reconstruction and scratch-training being 1.438 ± 0.140 (t(179) = 2.961, p = 7.726 ×10^−3^, two-sided t-test) and 1 (t(179) = 8.883, p = 2.228 ×10^−8^, two-sided t-test), respectively. The best performance across participants for these baseline methods were 1.600 ± 0.164 and 1, respectively. Detailed MOS scores for both natural and re-synthesized speech can be found in **supplementary Fig. 2**.

In summary, our results showed significant improvements in the quality of re-synthesized speech compared to other baseline models, reaching a level that was intelligible to humans. The re-synthesized speech obtained from different participants’ data exhibited similar consistent levels of intelligibility, indicating the generalizability and applicability of our approach across various participants.

### Temporal landmarks can be reliably detected from neural-driven re-synthesized speech

The previous section mainly concerns the acoustic quality of the neural-driven reconstructed speech. Furthermore, the perception of speech also involves detection and decoding of higher-levels of information, including phonemes, syllables, and words, within natural speech. The inquiry into whether humans can perceive these language units encompasses two aspects: firstly, the discernment of their positions within the re-synthesized speech audio, and secondly, the ability to decode these language units from re-synthesized speech audio signal. In this section, we introduce a series of binary classifiers aimed at investigating the detection of different temporal landmarks regarding phoneme, syllable, and word onsets from the neural-driven re-synthesized speech audio.

To probe the detection of various language units from the re-synthesized speech audio, we implemented the detectors of phoneme, syllable, and word onsets in the re-synthesized sentence-level speech, as illustrated in (**Fig. 3a-b**). The detection models were trained, validated on TIMIT and test on the neural-driven re-synthesized speech. The F1-scores for phoneme, syllable, and word onset detection were 86.3% ± 3.2%, 86.2% ± 2.9%, and 72.8% ± 2.6%, respectively. In comparison, the performances of the onset detectors for the actual original natural speech in the TIMIT test set yielded F1-scores of 93.9% ± 2.1%, 93.3% ± 5.0%, and 79.2% ± 9.6%, respectively (**Fig. 3c**). The detectors’ F1-scores for phoneme-level and syllable-level temporal landmarks in natural speech from TIMIT were significantly better than those for word-level landmarks (phoneme: t(179) = 15.109, p = 3.021 ×10^−18^, syllable: t(179) = 14.950, p = 1.322 ×10^−17^, two-sided t-test). The precision and recall for word onsets were 77.6% ± 3.9% and 71.5% ± 4.0%, respectively, whereas the performance for natural speech yielded precisions and recalls of 85.0% ± 7.3% and 75.9% ± 7.5% (**Fig. 3d**). The results revealed notable distinctions across different types of noise interference: phoneme, syllable and word onsets exhibited superior performance compared to babble noise at SNR=20dB (phoneme: t(198) =1.822, p = 0.070; syllable: t(198) = 1.883, p = 0.061; word: t(198) = 0.785, p = 0.433, two-sided t-test). Word onset detection also showed better performance on precision and recall than raw speech with babble noise at SNR=20dB (precision: t(198) = 0.512, p = 0.609, two-sided t-test; recall: t(198) = 0.968, p = 0.334, two-sided t-test).

**Fig 3.**
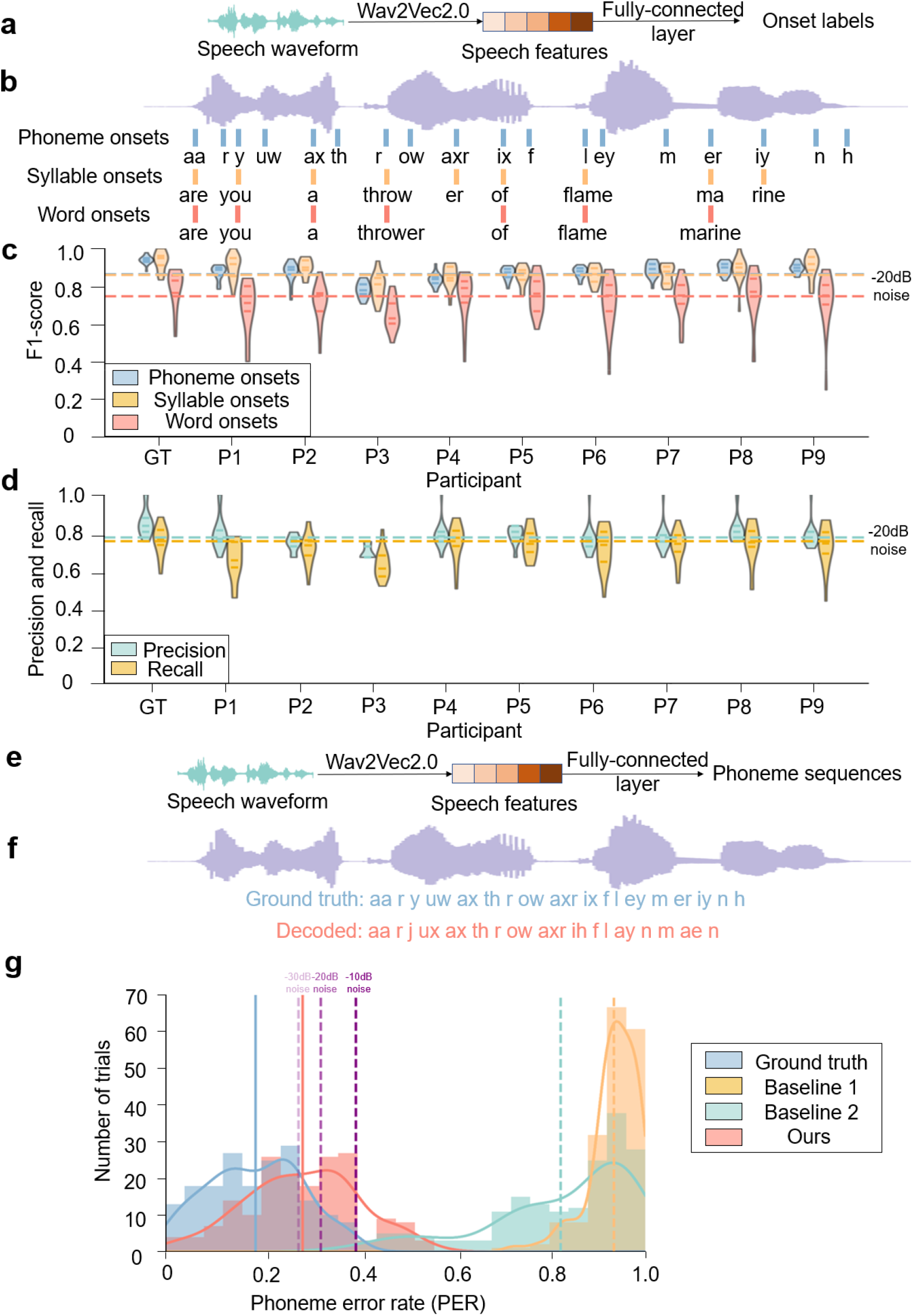
Evaluation of re-synthesized speech on temporal landmark detection and phoneme recognition. a.The framework illustrates the process of temporal landmark detection fine-tuned on Wav2Vec2.0, where linear binary classifiers are trained to discern whether fixed waveform segments contain the onset of phonemes, syllables, or words. b.An example of temporal landmark detection results, with blue, yellow, and green lines marking the detected onsets of phonemes, syllables, and words within a sentence, respectively. Underneath the waveform, annotations denote the corresponding phonemes, syllables, or words. c. The F1-scores summarize the performance of temporal landmark detection across participants for phonemes, syllables, and words, reflecting the model’s capability in accurately identifying these linguistic units. The F1-score of temporal landmark detection on speech with added -20dB babble noise is represented by corresponding colored dashed lines. GT: ground truth (raw speech). P1-P9: participant 1 – participant 9. d. An examination of precision and recall specifically for word onset detection across various participants, highlighting the model’s balance between correctly identifying word onsets and capturing all relevant instances. The precision and recall of word onset detection on speech with added -20dB babble noise is represented by corresponding colored dashed lines. The x-axis labels mean the same as those in figure c. e. Depicted is the methodology for automatic phoneme decoding, utilizing a Wav2Vec2.0-fine-tuned linear classifier to translate entire input speech sentences into phoneme sequences. f. An illustration showcases a decoded phoneme sequence, contrasting the grey waveform representing raw speech with the purple waveform symbolizing the synthesized speech output. g. Individual trial Transcription PER analyses for differing re-synthesis techniques are presented alongside their respective Kernel Density Estimation (KDE) curves, offering insights into the variability and overall quality of the speech synthesis process. The vertical lines represent the mean values of the corresponding groups of KDE curves. The purple vertical lines from light to dark illustrate the change in mean PER values with increasing levels of babble noise (-30, -20 and -10dB).

The differentiation in word onset detection mainly affects recall due to the subtle distinction between syllable and word onsets in speech waveforms. These similarity results in some syllable onsets being misidentified as word onsets when analyzing merely the speech waveform, causing a decrease in recall for word-level landmarks compared to syllable-level landmarks in natural speech from TIMIT. The consistency of temporal landmark detection results across different participants indicates the reliability of the findings.

### Phonemic information decoded from the reconstructed speech

After addressing the problem of detecting temporal landmarks, we proceeded to investigate whether the content of these linguistic units could be decoded from the neural-driven re-synthesized speech audio. To quantitatively evaluate the intelligibility of the re-synthesized speech, we developed an automatic phoneme recognizer to transcribe speech waveforms into phoneme sequences using natural speech sentences from TIMIT corpus (**Fig. 3e-f**).

This phoneme recognizer was fine-tuned based on a pre-trained Wav2Vec2.0 model[20]. Utilizing pre-trained Wav2Vec2.0 embedding features, a fully-connected phoneme classification layer transcribed the speech waveform into a phoneme sequence. The phoneme recognizer was trained, validated on TIMIT and tested on re-synthesized speech audio derived from the recorded neural activities. The phoneme recognizer achieved an averaged phoneme error rate (PER) of 28.6% ± 11.0% (mean ± s.d.) on the neural-driven re-synthesized speech across all 9 participants (Fig. 3g), with the best performance across participants being 26.1% ± 10.3%. As a benchmark, the phoneme recognizer achieved an averaged PER of 18.9% ± 8.2% on the original raw speech. Compared to the reconstructed speech from other baseline models, our neural-driven re-synthesized speech closely resembled the ground truth raw speech in terms of the PER distribution (**Fig. 3g**). Importantly, our neural-driven re-synthesized speech outperform both baseline models (MLP-reconstructed speech: PER = 82.3% ± 15.5%, t(179) = 19.875 and p = 1.123 ×10^−21^ for two-sided t-test; scratch-training speech: PER = 93.4% ± 5.1%, t(179) = 42.414, p = 1.293 ×10^−33^, two-sided t-test). In terms of PER, the quality of the reconstructed speech outperforms the raw speech waveform with -20dB additive noise, and was comparable to that with -30dB additive noise (SNR = 20dB: t(198) = 3.656, p = 7.724 ×10^−4^,two-sided t-test; SNR=30dB: t(198) = -0.286, p = 0.775, two-sided t-test).

### Subjective word perception of the re-synthesized speech by human evaluators

Finally, we examined the subjective recognition of words by human evaluators as a quantitative evaluation of the higher-level speech reconstruction intelligibility. The assessment involved four individuals who listened to the original speech and the re-synthesized speech derived from ECoG recordings in a randomized order. After each sentence, the evaluators were asked to transcribed the sentence on a word-by-word basis, and the word recognition accuracy (WRA) for each sentence were calculated. The evaluators successfully recognized 71.6% of all the words in the neural-driven reconstructed speech from the test set (**Fig. 4a)**. Notably, the highest performance across participants was 79.5%, while the best performance across evaluators was 75.78%. The accuracy was consistent with the previously computed objective phoneme error rate, providing converging evidence that our proposed method could reliably reconstruct intelligible speech.

**Fig 4.**
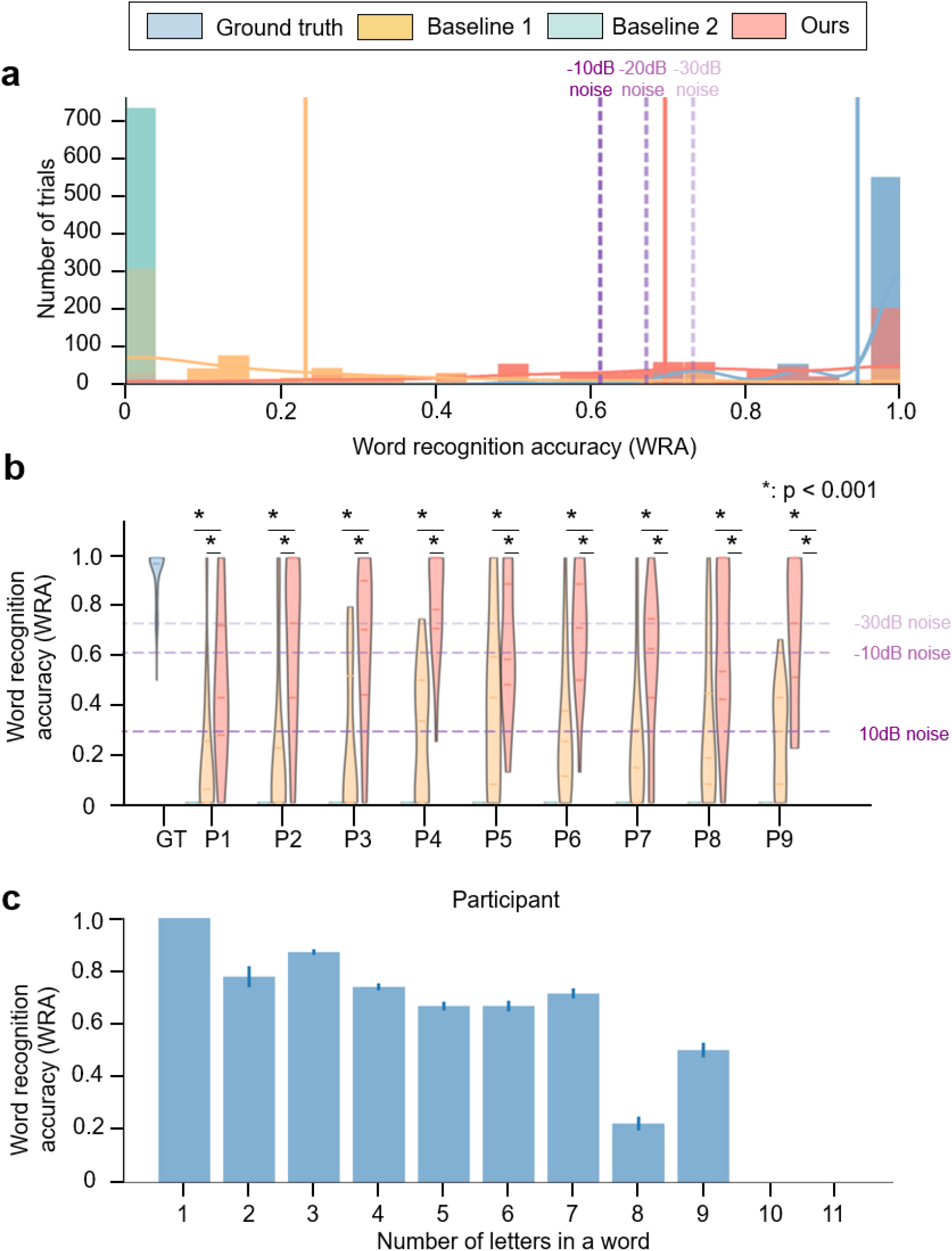
Word Recognition Accuracy (WRA) of re-synthesized speech. a.Transcription Word Recognition Accuracy (WRA) for separate trials under various re-synthesis methodologies, accompanied by their respective Kernel Density Estimation (KDE) curves. This depiction offers insights into the distribution of recognition performance across trials, further emphasizing the 71.6% overall word recognition achieved, with standout individual performance peaking at 79.5%. The purple vertical lines from light to dark illustrate the change in mean WRA values with increasing levels of babble noise (-30, -20 and -10dB). b.WRA of re-synthesized speech for each participant across different re-synthesis techniques, aggregating to an average of 68.8% ± 6.1%. This comprehensive view captures the best performance across participants at 81.0% ± 4.5%, reinforcing the method’s consistent intelligibility and generalizability across a diverse set of listeners. The three shades of purple dashed lines from light to dark represent the average WRA after adding babble noise at 10 dB, -10 dB, and -30 dB to the original speech waveform. c.Analysis of WRA with respect to the word length, explicitly showing that words consisting of seven or fewer letters achieve a significantly higher recognition rate of 76.9%, in comparison to words exceeding this length, which are recognized at a notably lower rate of 23.1%. This segment underscores the impact of word length on recognition efficacy, deepening our understanding of the human being’s perception of natural speech.

The WRA across all re-synthesized speech sentences from all 9 participants is 68.8% ± 6.1%, with the best performance across participants at 81.0% ± 4.5% and the best performance across evaluators at 77.4% ± 4.3% (**Fig. 4b**). In contrast, other baseline methods did not achieve high intelligibility: scratch-trained speech yielded a WRA of 0.0%, and the MLP-reconstructed speech achieved a WRA of 23.3% ± 6.5% (MLP-reconstructed speech: t(719) = 15.190, p = 1.976 ×10^−-27^, two-sided t-test; scratch-trained speech: t(719) = 66.448, p = 3.793 ×10^−309^,two-sided t-test). These findings demonstrate that our re-synthesized speech exhibits good intelligibility at the word level. These results indicate that our re-synthesis method produces speech with word-level intelligibility that is consistent across different participants, demonstrating its generalizability.

Additionally, we found that longer words tended to have lower WRA, compared to shorter words. Notably, words less than seven letters had a significantly higher WRA (76.9%) compared to words with more than seven letters (23.1%, t(4822) = 13.031, p = 5.329 ×10^−15^, two-sided t-test) (**Fig. 4c**). Detailed evaluation results across the four recruited evaluators were provided in **Supplementary Fig. 4**.

## Discussion

In this study, we present a novel approach to re-synthesize high-fidelity natural speech from electrocorticography (ECoG) signals using a pre-trained encoder-decoder framework. The key finding is that the generative speech model, employing a context-dependent speech encoder and a generative adversarial network (GAN), effectively re-synthesizes natural speech from limited neural data. Specifically, using merely 20 minutes of ECoG recordings, the model achieved a phoneme error rate (PER) of 28.6% (**Fig. 3g**), while human listeners recognizing 71.6% of the words accurately (**Fig. 4a**). This combination of pre-trained models (Wav2Vec2.0 and HiFi-GAN) and feature alignment allows for a lightweight yet effective neural feature adaptor that translates ECoG signals into the latent space of a deep speech encoder. Such an approach reduces the need for extensive neural training data, traditionally a bottleneck in neural decoding. The results demonstrate the feasibility of integrating pre-trained deep speech models into neural decoding, paving the way for more advanced brain-computer interfaces and neuroprosthetics. These findings are supported by the analysis of word recognition accuracy (WRA) across participants, which averaged 68.8% ± 6.1%, peaking at 81.0% ± 4.5% under optimal conditions (**Fig. 4a, Fig. 4b, Supplementary Fig. 4**). Additionally, the study reveals that shorter words (seven or fewer letters) are recognized at a significantly higher rate (76.9%) compared to longer ones (23.1%), suggesting the impact of word length on recognition efficacy (**Fig. 4c**).

The approach in this work for speech reconstruction improves performance and data efficiency compared to previous methods by overcoming limitations in spatiotemporal resolution and using invasive ECoG recordings and advanced nonlinear models with pretraining. Our approach diverges from previous methods that focused on direct decoding from neural signals, which often faced challenges due to the demanding spatiotemporal resolution requirements of neural recordings. While end-to-end re-synthesis is intuitively appealing, prior related work has shown that direct re-synthesis methods exhibit modest performance due to the scarcity of available neural data[8]. To address these limitations, our study introduces a pre-trained encoder-decoder framework utilizing a nonlinear model and pretraining techniques. By employing a pre-trained generative adversarial network (GAN) and neural feature alignment, we achieve significant improvements in re-synthesis quality with invasive ECoG recordings. This method notably reduces the amount of neural data needed for effective speech decoding. Our model demonstrates promising performance using only 20 minutes of neural data, offering a more data-efficient solution for neural-to-speech decoding, rather than using hours[33] or even days[9] of neural recordings. Our approach further distinguishes itself by decoding entire sentences rather than discrete phonemes, syllables, or words, facilitated by high spatiotemporal resolution neural recordings and a pre-trained deep speech encoder and generator. Moreover, our proposed model combined a shared pre-trained generative model and participant-specific projection module, facilitating transferability across participants with minimal additional single-participant level training. This advancement addresses the limitations of direct decoding techniques and enables improved re-synthesis quality and more efficient use of neural data.

One of the key roles of speech decoding is to advance our understanding of neural coding by assessing the quality of synthesized speech. Our work enhances this by enabling a comprehensive evaluation at the sentence level, incorporating metrics beyond ESTOI (**Fig. 2b**), mel-spectrogram MSE (**Fig. 2c**), MOS (**Fig. 2d, Supplementary Fig. 2**), etc., which were not fully addressed in previous studies[7, 8, 18]. We conducted a multi-level analysis of synthesized speech, examining various language units from low to high levels, including temporal landmarks detection for phonemes, syllables, and words, automatic phoneme recognition, and human evaluation of re-synthesized sentences. Our results show that most temporal landmarks can be detected, approaching machine limits, over 70% of phonemes can be recognized with a simple automatic recognizer (**Fig. 3g**), and human evaluators can identify approximately 70% of words (**Fig. 4a, Fig. 4b, Supplementary Fig. 4**).

The findings reinforce that pre-trained AI models can effectively align with neural representations, bridging brain signals and speech, supporting the development of more generalizable brain-to-AI interfaces and improving neural decoding through advanced generative models. Related works[22, 24] has demonstrated the representational similarity between neural activity and AI model outputs, indicating that self-supervised speech models can successfully link brain signals with speech representations. This alignment signifies a significant step towards brain-AI integration, where the discovery of simpler mappings between neural and AI model spaces can facilitate practical applications, such as speech synthesis and cognitive interfaces. Moreover, our findings support the notion that deep learning models, pre-trained on extensive datasets, can serve as “cognitive mirrors,” reflecting the intricate processes by which the human brain handles speech and other high-level functions [34]. With the advent of more sophisticated generative models, we anticipate enhanced neural decoding capabilities, including the potential to improve signal quality (SNR) and refine the generative model itself. Additionally, this framework extends beyond speech to other modalities, such as vision, suggesting that similar principles may apply to the generation of visual content from neural signals.

This work advances speech neuroprosthetics and BCIs by enabling real-time speech reconstruction with minimal neural data, improving scalability and practicality through pre-trained networks, and suggesting potential for broader applications in perception, memory, and emotion. By adapting neural activity to a common latent space of the pre-trained speech Autoencoder, our framework significantly reduces the quantity of neural data required for effective speech decoding, thereby addressing a major limitation in current BCI technologies. This reduction in data needs paves the way for more accessible and scalable BCI solutions, particularly for individuals with speech impairments who stand to benefit from immediate and intelligible speech reconstruction. Furthermore, the applicability of our model extends beyond speech, hinting at the possibility of decoding other cognitive functions once the corresponding neural correlates are identified. This opens up exciting avenues for expanding BCI functionality into areas such as perception, memory, and emotional expression, thereby enhancing the overall quality of life for users of neuroprosthetic devices.

There are several limitations in our study. The quality of the re-synthesized speech heavily relies on the performance of the generative model, indicating that future work should focus on refining and enhancing these models. Currently, our study utilized English speech sentences as input stimuli, and the performance of the system in other languages remains to be evaluated. Regarding signal modality and experimental methods, the clinical setting restricts us to collecting data during brief periods of intraoperative wakefulness, which limits the amount of usable neural activity recordings. Overcoming this time constraint could facilitate the acquisition of larger datasets, thereby contributing to the re-synthesis of higher quality natural speech. Additionally, exploring non-invasive methods represents another frontier; with the accumulation of more data and the development of more powerful generative models, it may become feasible to achieve effective non-invasive neural decoding for speech re-synthesis.

In summary, we introduce a novel approach using a pre-trained encoder-decoder framework to re-synthesize high-fidelity sentence-level natural speech from cortical recordings, demonstrating the effectiveness of this framework. Despite the challenges associated with the performance reliance on generative models and the limitations imposed by the intraoperative awake period for data collection, the research lays a solid foundation for future advancements. Moving forward, refining these models and exploring non-invasive methods will be key areas of focus. The integration of deep neural networks with neurophysiological data holds significant promise for advancing our theoretical understanding of speech processing in the brain, as well as enhancing practical applications of brain-computer interfaces, potentially revolutionizing the field of neuroprosthetics and improving the lives of individuals reliant on such technologies. This advancement paves the way for broader cognitive functions exploration and promises transformative improvements in the functionality of neural prosthetic devices. Ultimately, this convergence of technologies will deepen our insights into how the brain processes speech and open new avenues for BCI applications.

## Methods

### Participants

The study comprised 9 monolingual right-handed participants, each with electrodes placed on the left hemisphere cortex to clinically monitor seizure activities. Electrode placement followed specific clinical requirements (see **Supplementary Fig. 1** for grid placement in each placement). Participants were thoroughly informed about the experiment, as outlined in the consent document approved by the Institutional Review Board (IRB) of the University of California, San Francisco. They volunteered for participation, ensuring that their involvement did not influence their clinical care. Additionally, verbal consent was obtained from participants at the beginning of each experimental session.

### Data acquisition and neural signal processing

All participants utilized high-density ECoG grids of uniform specifications and type. During the experimental tasks, neural signals captured by the ECoG grids were acquired using a multi-channel amplifier optically linked to the digital signal processor. Data recording was performed through Tucker-Davis Technology (TDT) OpenEx software. The local field potential at each electrode contact was amplified and sampled at 3052 Hz. Subsequent to data collection, the experiment employed the Hilbert transform to compute the analytic amplitudes of eight Gaussian filters (with center frequencies ranging from 70 to 150 Hz), and the signal was down-sampled to 50 Hz. These tasks were segmented into record blocks lasting approximately 5 minutes each. The neural signals were z-score normalized for each recording block.

### Experimental Stimuli

The acoustic stimuli employed in this study comprised natural and continuous English speech. The English speech stimuli were sourced from the TIMIT corpus, consisting of 499 sentences read by 286 male and 116 female speakers. A silence interval of 0.4 seconds was maintained between sentences. The task was organized into five blocks, each with a duration of approximately 5 minutes.

### Electrode localization

In cases involving chronic monitoring, the electrode placement procedure involved pre-implantation MRI and post-implantation CT scans. In awake cases, temporary high-density electrode grids were utilized to capture cortical local potentials, with their positions recorded using the Medtronic neuronavigation system. The recorded positions were then aligned to pre-surgery MRI data, with intraoperative photographs serving as additional references. Localization of the remaining electrodes was achieved through interpolation and extrapolation techniques.

### Speech-responsive electrodes selection

The selection of speech-responsive electrodes played a crucial role in the re-synthesis of high-quality natural speech and avoid overfitting. Onsets were identified as the initiation of speech and are preceded by more than 400 ms of silence. A paired sample t-test was conducted to investigate whether the average post-onset response (400 ms to 600 ms) exhibited a significant increase compared to the average pre-onset response (-200 ms to 0 ms, p < 0.01, one-sided, Bonferroni corrected).

### Architecture and training of the speech Autoencoder

The speech Autoencoder served as a pivotal component, taking natural speech as input and reproducing the input itself. Within this model, the intermediate encoding layers were construed as the feature extraction layers. The architecture comprises an encoder, namely Wav2Vec2.0, and a decoder, which is the HiFi-GAN generator.

The Wav2Vec2.0 encoder was comprised of 7 1D convolutional layers that down-sample 16kHz natural speech to 50Hz, thereby extracting 768-dimensional local features. Additionally, it integrated 12 transformer-encoder blocks to extract contextual feature representations from unlabeled speech data.

On the other hand, the HiFi-GAN generator incorporated 7 transposed convolutional layers for up-sampling, along with multi-receptive field (MRF) fusion modules. Each MRF module aggregated the output features from multiple ResBlocks. The discriminator within the HiFi-GAN encompassed both multi-scale and multi-period discriminators. Notably, the up-sample rates and kernel sizes of each ResBlock in the MRFs were meticulously configured. The up-sample rates of each ResBlock in the MRFs were set as 2, 2, 2, 2, 2, 2, 5 respectively. The up-sample kernel sizes of each ResBlock in the MRFs were set as 2, 2, 3, 3, 3, 3, 10. The periods of the multi-period discriminator were set as 2, 3, 5, 7, 11.

The HiFi-GAN generator *G* and discriminators *D*_*k*_ were trained simultaneously and adversarially. The loss function consisted of three parts: adversarial loss *L*_*Adv*_ to train the HiFi-GAN, mel-spectrogram loss *L*_*Mel*_ to improve the training efficiency of the generator and feature matching loss *L*_*FM*_ to measure the similarity between original speech and re-synthesized speech from learned features:

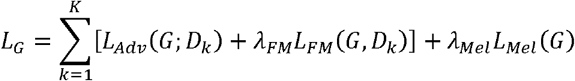

The regularization coefficients *λ*_*FM*_and *λ*_*Mel*_ were set as 2 and 50 respectively in the implementation. In detail, the three terms of *L*_*G*_ were:

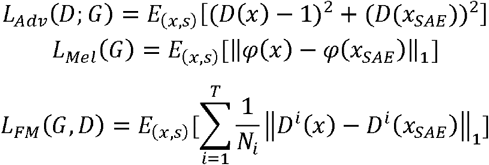

Where *φ* represented the mel-spectrogram operator, *T* was the number of the layers in the discriminator, *D*^*i*^ and *N*_*i*_ were the features and the number of features in *i* th layer of the discriminator, *x* and *x*_*SAE*_ represented the original speech and the reconstructed speech using the pre-trained speech Autoencoder:

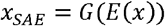

The parameters in Wav2Vec2.0 were frozen within this training phase. The parameters in HiFi-GAN were optimized using Adam optimizer with a fixed learning rate as 10^−5^, β_*1*_ = 0.9, β_*2*_ = 0.999. We trained this Autoencoder in LibriSpeech, a 960-hour English speech corpus with a sampling rate of 16kHz. We spent 12 days parallelly train on 6 Nvidia GeForce RTX3090. The maximum training epoch was 2000. The optimization did not stop until the validation loss no longer decreased.

### Mean opinion score (MOS) and extended short time objective intelligibility (ESTOI)

The Mean Opinion Score (MOS) is a subjective metric employed to assess the intelligibility of synthesized speech. In order to obtain MOS ratings, we recruited a cohort of 20 volunteers who possessed a minimum of 10 years of English learning experience. These individuals were tasked with evaluating the quality of the synthesized speech produced by the pre-trained speech Autoencoder. The evaluation was conducted using a numerical scale ranging from 1 (bad) to 5 (excellent). To ensure the validity and reliability of the ratings, the following criteria were upheld: (1) participants had a proficiency in speaking English for over 10 years; (2) participants were not involved in any aspects of the research work apart from the MOS scoring; (3) there were no conflicts of interest between the participants and the researchers; (4) participants were neither patients, family members of patients, nor employees of hospitals.

In addition to MOS, we utilized the Extended Short-Time Objective Intelligibility (ESTOI) score as an objective indicator to assess the intelligibility of the speech synthesis technology. The ESTOI score ranges from 0 (worst) to 1 (best). This metric captures distortions in the spectrotemporal modulation patterns of speech and is particularly sensitive to inconsistencies in spectral and temporal patterns.

### Babble noise

Babble noise is a specific type of noise originating from multiple speakers conversing simultaneously, typically experienced in public places such as restaurants, conference halls, or busy streets. This noise presents a challenge to speech processing systems primarily because it exhibits speech-like characteristics—it contains natural fluctuations and spectral properties similar to human speech. The similarity in the frequency domain increases the difficulty of speech recognition and understanding.

Compared to white noise, babble noise has several notable differences. Firstly, white noise has uniform energy distribution across all frequencies, whereas babble noise concentrates higher energy in certain frequency regions (such as where speech resonances occur) due to its composition of human speech. Secondly, the temporal characteristics of babble noise are non-stationary, changing over time, whereas white noise is stationary with characteristics that do not vary over time. Thirdly, due to its speech-like nature, babble noise prompts the human ear to attempt deciphering speech content, making it more challenging for speech recognition systems as it blurs the boundary between background noise and target speech. Lastly, generating babble noise typically involves considerations of factors like the number of speakers, conversation topics, and emotions, whereas white noise is an idealized, featureless noise model.

To synthesize babble noise, we randomly selected sentences from the training set of the TIMIT corpus as interference signals. Specifically, these sentences came from different speakers, sampled from their recorded conversation segments. Subsequently, these segments were overlapped randomly to simulate real-world conversational scenarios. During the overlapping process, considerations such as turn-taking between speakers, relative strength and direction of each speaker’s voice were typically considered to enhance the realism of the noise. The resulting synthesized babble noise was a multi-channel mixture containing the effect of multiple speakers conversing simultaneously.

### Neural feature adaptor training and ablation test

The speech re-synthesize model was trained to finally acquire the re-synthesized natural speech. The speech re-synthesize model received *z*-scored high gamma ECoG snippets with dimensions *N* × *T*, where *N* was the number of selected speech-responsive electrodes, and *T* was the ECoG recordings (50 Hz) down-sampled to the same sample rate as the speech features extracted from pre-trained Wav2Vec2.0.

A three-layer bidirectional LSTM adaptor *f* (see Fig. 1 D) aligned recorded ECoG signals to speech features. The output of the adaptor was *768* × *T*, as the input of the HiFi-GAN generator pre-trained in phase 1. The ultimate output of the HiFi-GAN was a speech waveform with a length of 1 × 320*T* + 80) in a 16K sample rate.

We froze all the parameters of the pre-trained speech Autoencoder while training the adaptor. The loss function used in this phase consisted of two parts: the mel-spectrogram loss *L*_*Mel*_ and the Laplacian loss *L*_*Lap*_:

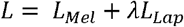

Where the regularization coefficient λ was set as 0.1 using the ablation test.

*L*_*Mel*_ was used to enhance the acoustic fidelity and intelligibility of the ultimate re-synthesized speech from recorded neural activity:

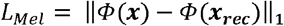

where *x* and *x*_*rec*_ respectively represented the original and re-synthesized speech, *Ф* represented the mel-spectrogram operator.

In order to improve the phonetic intelligibility, a Laplace operator as a convolution kernel was operated to the mel-spectrogram, which was a simplified representation of the convolution on the spectrogram, while convolution on spectrogram was proven to be effective on speech denoising and enhancement[35]:

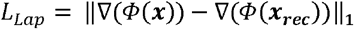

The parameters in HiFi-GAN generator were frozen within this training phase. The parameters in the adaptor was trained in Stochastic Gradient Descent (SGD) optimizer with a fixed learning rate of 3 × 10^−3^ and a momentum of 0.5. A 10% dropout layer was added in this stage to avoid overfitting. The optimization did not stop until the validation loss no longer decreased.

We respectively selected uni-directional, bi-directional LSTM layers and Transformer blocks as the adaptor to adapt ECoG recordings to speech feature layers. **Supplementary Fig. 3** represents the ESTOI score using LSTM layers (ranges from l layer to 5 layers, both uni-directional and bi-directional are included) and Transformer blocks (ranges from 1 block to 5 blocks) as the adaptor. We used from 128 to 4096 dimensional (2 as the log scale) hidden size in the LSTM layers. The number of attention heads and the dimension of the hidden space were set as 8 and 80 in the Transformer block.

For each participant, we trained a personalized feature adaptor. Each individual feature adapter underwent training for 500 epochs on one NVidia Geforce RTX3090 GPU, which required 12 hours to complete. The dataset was divided into training, validation, and test sets, comprising 70%, 20%, and 10% of the total trials, respectively.

### Two baselines used in re-synthesized speech evaluation

The re-synthesized speech sentences obtained from recorded neural activity are assessed using two end-to-end speech re-synthesis methods as our baselines (**Fig. 2a**). The first baseline model employed a three-layer multi-layer perceptron (MLP) as a regression model to directly map neural recordings onto the mel-spectrogram without any pre-trained decoder. This baseline was designed to highlight the limited quality of direct end-to-end synthesizing natural speech from recorded neural activities. As for the second baseline, we trained a speech re-synthesis model from scratch without Librespeech pre-training, employing the same architecture depicted in **Fig. 1**, In this baseline, all parameters in the adaptor and HiFi-GAN were made tunable, rather than solely tuning the adaptor. This baseline aimed to demonstrate the importance of the pre-training stage and to evaluate the extent to which the re-synthesized speech outperforms training from scratch when incorporating the pre-training phase of the deep speech generator.

### Onset detection of re-synthesized speech sentences

The onset detectors in our study functioned as binary classifiers that determine the existence of phoneme, syllable, or word onsets within speech periods. To accomplish this, the classifiers were provided with fixed-length segments of speech waveforms (0.04s for phonemes and 0.2s for syllables and words). The output of these classifiers was a scalar value ranging from 0 to 1, representing the probability of the existence of a temporal landmark. Each classifier comprises a Wav2Vec2.0 feature extractor and a logistic regression-based classifier, which includes a linear layer and a sigmoid layer.

To address the challenge posed by the significant disparity in the quantity of positive and negative samples, data augmentation techniques were applied to the speech segments during training. These techniques included volume augmentation, speed changes, pitch shifts, and the addition of Gaussian noise, among others. By employing these augmentation methods, we aimed to mitigate any potential bias resulting from the imbalanced dataset. During training, both positive and negative speech segments were sampled at a quantity of 5000 each for the onset detector.

In order to assess the performance of the models, they were trained and evaluated using the TIMIT training set and TIMIT test set. Additionally, the models were tested on re-synthesized speech ECoG recordings. These models are trained using AdamW optimizer with a fixed learning rate of 0.0001, *β*_1_ = 0.9, *β*_2_ = 0.999, and *ε* = 10^−8^ using a cross-entropy loss function. The optimization did not stop until the validation loss no longer decreased.

### Automatic phoneme recognition of re-synthesized speech sentences

In order to quantitatively evaluate the synthesized speech is to recognize the phonemes, we trained a phoneme recognizer using Wav2Vec2.0, a logistic regression phoneme classifier, and a phoneme decoder. Similar to onset detection, Wav2Vec2.0 was used for feature extraction, the classifier outputs a probability vector from Wav2Vec2.0 features, and the phoneme decoder decoded a phoneme sequence from the probability vectors to finally get the phoneme sequence.

The models are trained and evaluated on the training set and test set of TIMIT, and tested on the re-synthesized from ECoG recordings. These models were trained using AdamW optimizer with a fixed learning rate of 0.0001, *β*_1_ = 0.9, *β*_2_ = 0.999, and *ε* = 10^−8^ in CTC loss function. The optimization did not stop until the validation loss no longer decreased.

### Phoneme error rate (PER) and word recognition accuracy (WRA)

Phoneme error rate (PER) measures the degree of difference between the output of a phoneme transcriber and a reference transcription when transcribing speech into phonemes. PER is calculated by counting the total number of insertions, deletions, and substitutions required to align the Automatic Speech Recognition (ASR) system output with the reference transcription. The process of computing WER entails aligning the ASR system output with the reference transcription, counting the number of mismatched words, and dividing it by the total number of words in the reference transcription to obtain the error rate as a percentage.

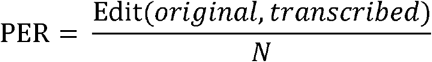

Where Edit(*original, transcribed*) denotes the edit distance between original and transcribed phoneme sequences; *N* is the number of phonemes in the sentence

Word recognition accuracy (WRA) is defined as the ratio of the number of the corrected recognized words to the total number of words. WRA is calculated as follows:

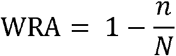

Where *n* is the number of correctly recognized words; *N* is the number of words in the sentence.

### Acquisition of WRA

We enlisted four individuals to assess the word-level quality of speech. These evaluators listened to original speech and re-synthesized speech derived from ECoG recordings, presented in a randomized order. After each sentence, we provided them with its transcription and asked them to report their recognition of the sentence on a word-by-word basis. This methodology enabled us to analyze the reconstructed natural speech at the word level. Specifically, it allowed for the calculation of Word Recognition Accuracy (WRA) for each sentence and facilitated observations of word recognition accuracy in relation to word length.

## Data availability

The raw datasets supporting the current study have not been deposited in a public repository because it contains personally identifiable patient information, but are available in an anonymized form from E.F.C upon reasonable requests.

## Code availability

All original code to replicate the main findings of this study can be found at https://github.com/CCTN-BCI/Neural2Speech. Any additional information required to reanalyze the data reported in this paper is available from the lead contact upon request.

## Competing interests

The authors declare no competing interests.

## Acknowledgments

This work is supported by the National Natural Science Foundation of China (32371154, Y.L.), Shanghai Rising-Star Program (24QA2705500, Y.L.), Shanghai Pujiang Program (22PJ1410500, Y.L.), and the Lingang Laboratory (LG-GG-202402-06, Y.L.). The computations in this research are supported by the HPC Platform of ShanghaiTech University.

## Notes

### Competing Interest Statement

The authors have declared no competing interest.

