## Supplementary Information for "Natural speech re-synthesis from direct cortical recordings using a pre-trained encoder-decoder framework"

Li *et al*

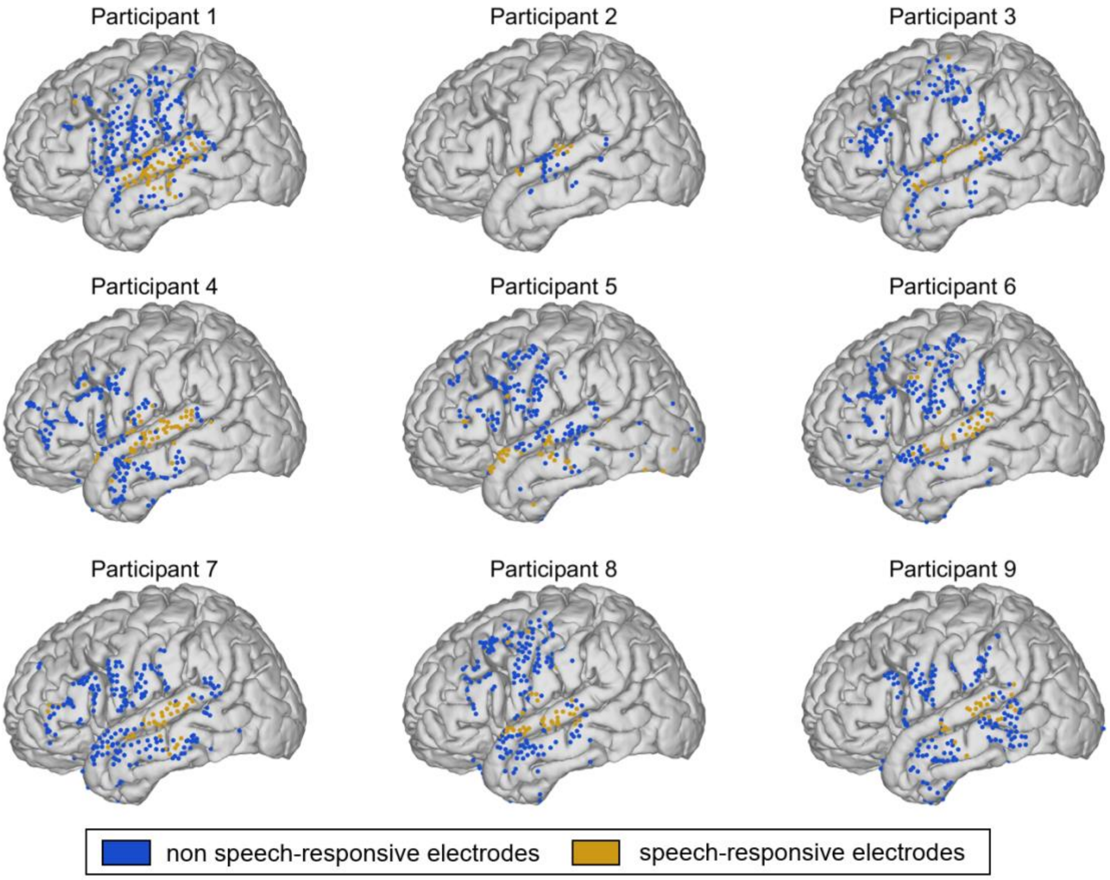


**Supplementary Figure 1. Speech responsive and tone discriminating electrodes for all participants.** ECoG grids covering the lateral temporal lobe of all participants were warped onto the MNI152 template. Yellow electrodes are responsive to speech, while blue electrodes are not.

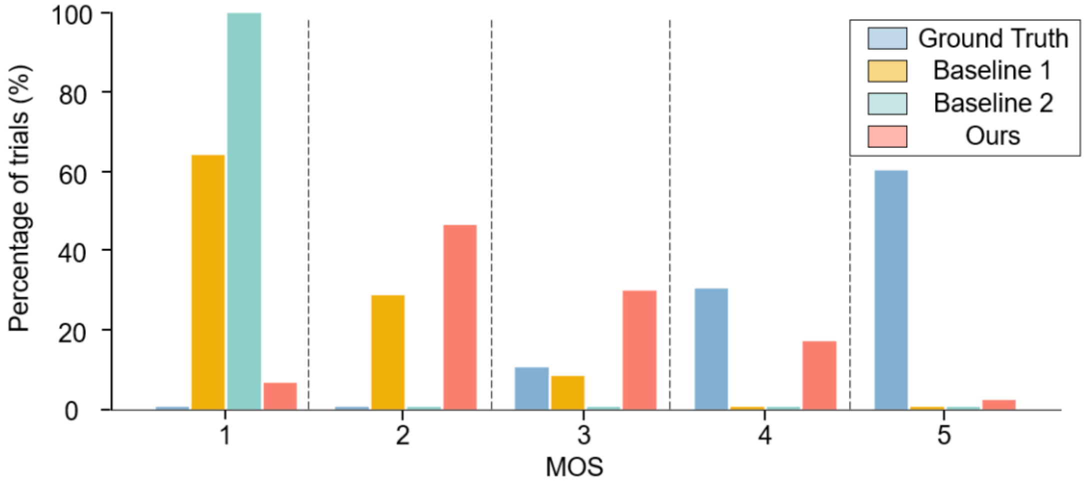


**Supplementary Figure 2 Detailed mean opinion score (MOS) of natural and re-synthesized speech, showing the percentage of trials for which human evaluators gave each MOS rating from worst (1) to best (5).** The x-axis shows the MOS rating, while the y-axis shows the percentage of trials in which that rating was given.

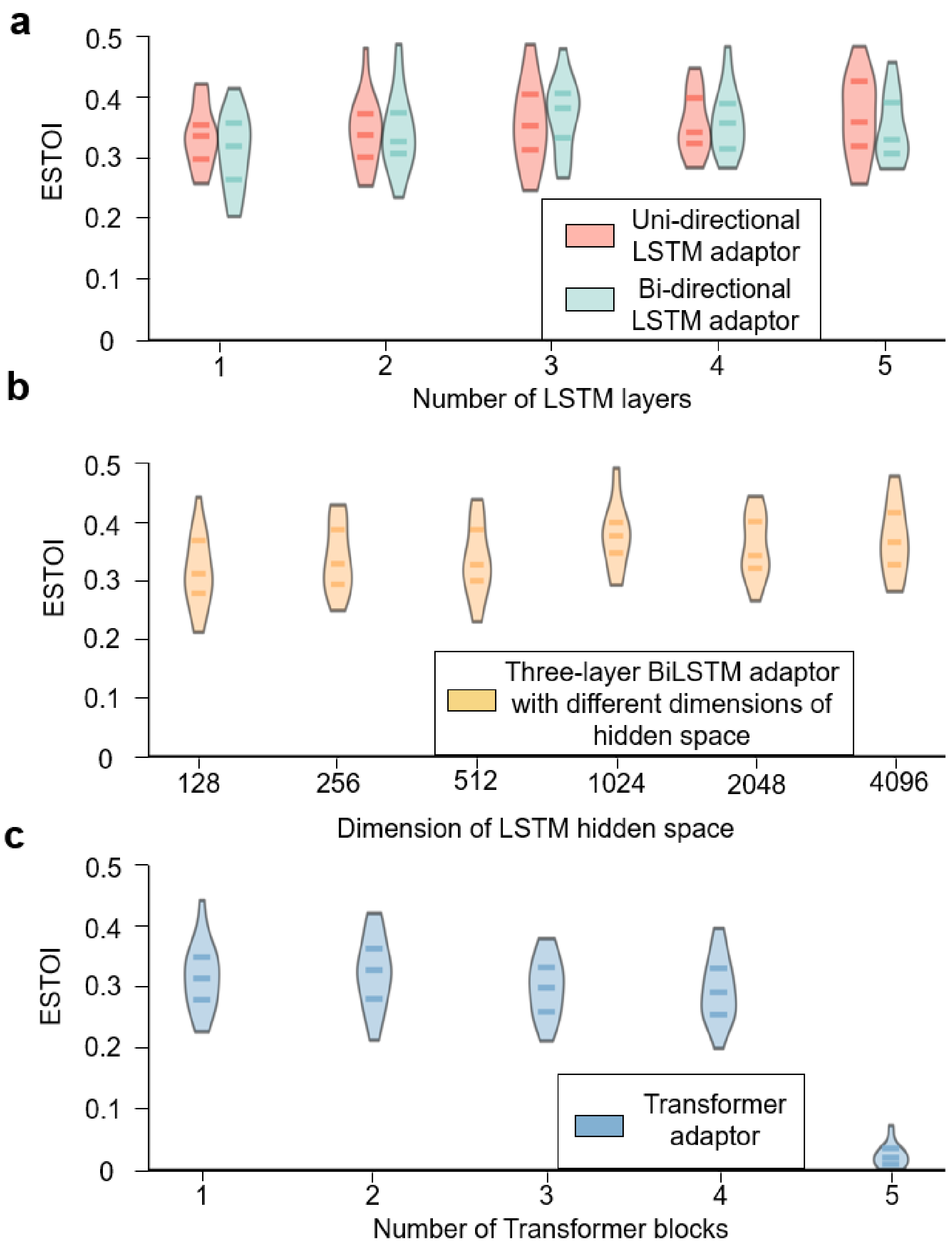


**Supplementary Figure 3. Ablation test of the adaptor.** In panel (a), the system's performance is compared when using either a uni-directional or bi-directional LSTM as the adaptor, while also varying the number of LSTM layers from 1 to 5. In panel (b), the impact of changing the dimensionality of the hidden space within the LSTM is examined, with values spanning a logspace from 126 to 4096. Finally, panel (c) compares the system's performance when using a Transformer instead of an LSTM as the adaptor. Overall, these results suggest that the choice of adaptor can have a significant impact on the quality of the re-synthesized speech.

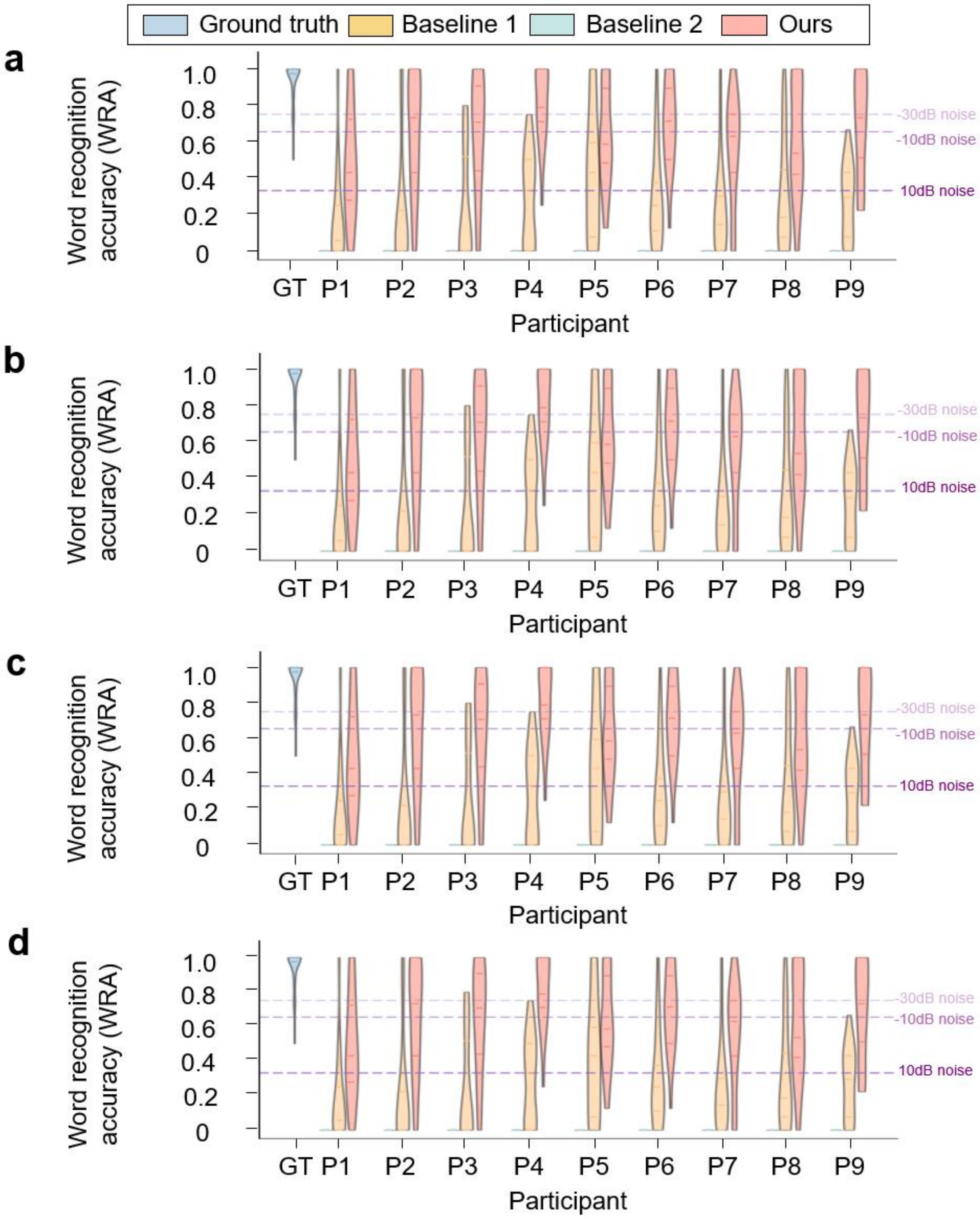


**Supplementary Figure 4. Word Recognition Accuracy (WRA) of re-synthesized speech across every one of the four evaluators, comparing against the ground truth and two baseline methods.** Each panel represents the WRA for a specific participant, with the purple lines indicating the WRA for the ground truth speech at various noise levels (-30dB, -10dB, and 10dB).
